# N-Glycan Fingerprinting of the NIST monoclonal antibody (NISTmAb)

**DOI:** 10.1101/2025.02.15.638459

**Authors:** Zhengliang L Wu

**Affiliations:** Bio-orthogonal Therapeutics, Edina, MN 55436, USA

**Keywords:** Nist mAb, Humira®, glycan ladder

## Abstract

Glycosylation on therapeutic antibodies is critically important for their drug efficacies. Here, using NISTmAb and Humira® as examples, we present methods of glycan fingerprinting for these antibodies. Glycans are first released with PNGase F or Endo S2, and then labeled by specific glycosyltransferases, including sialyltransferase ST6Gal1, fucosyltransferase FUT9, N-acetyl-glucosaminyltransferase MGAT3 and fucosyltransferase FUT8, with respective fluorophore-conjugated donor sugars. The labeled glycans are then separated by gel electrophoresis (SDS-PAGE). A fluorophore-labeled hyaluronan ladder is run along with the samples to reveal the relative mobility of each glycan band. Pretreatment of the samples with specific glycosidase or glycosyltransferase results in additional mobility shift of specific glycan bands, which allows identification of some of these bands. Particularly, we report the identification of α-Gal epitopes, core-6 fucosylated glycans, paucimannose glycans and likely some bisecting/tri-antennary glycans on NISTmAb. Overall, our methods could serve as quick assessments of glycosylation on therapeutic antibodies.

## Introduction

Glycosylation is a common post-translational modification and plays roles in protein stability, solubility, and affect cell-cell interactions [1, 2]. Therapeutic antibodies are the highest earning category of all biological drugs [3]. Glycosylation on the Fc region of therapeutic antibodies greatly affects their drug efficacies and safety [4, 5]. Especially, galactosylation, fucosylation and sialylation are well-established factors to effect IgG functions, ranging from inhibitory/anti-inflammatory efficacies to complement activation and antibody-dependent cellular cytotoxicity [6]. For these reasons, glycan analysis becomes critically important for therapeutic antibody drug development and production.

However, glycan analysis on therapeutic antibodies have been challenging due to lack of consistent and quantitative methodology, which is exemplified by the interlaboratory study on the glycosylation of NISTmAb, a humanized IgG1k monoclonal antibody produced in NS0 cells that has been used as a standard for testing *de novo* sequencing algorithm on bottom-up mass spectrometry [7]. The samples were analyzed by 66 labs around the world using various chromatography and mass spectrometry methods for glycan separation and identification and the diversity of the results was large with the number of glycan compositions identified by each laboratory ranging from 4 to 48 [8]. Among these labs, 32 reported galactosylated glycans, 37 reported core fucosylated glycans, and 13 reported α-Gal epitopes containing glycans.

Previously we reported glycan fingerprinting on Covid-19 RBD proteins with fluorophore-labeled sialic acid and fucose by ST6Gal1 and FUT8, respectively [9]. A challenge of this technique is to establish a glycan ladder so that any glycan under investigation can be referenced and compared. Here we report glycan fingerprinting on NISTmAb and the therapeutic antibody Humira® [10] using a fluorophore-labeled hyaluronan (HA) based glycan ladder [11]. The HA ladder contains more bands than our previous glycan ladder and allows us to assign precise relative mobility to any resolved glycan in a gel. In addition, we introduce FUT9 and MGAT3 as labeling enzymes in this report. Compared to ST6Gal1 that recognizes only terminal lactosamine (LN) structure [12], FUT9 also recognizes internal LN structures [13], therefore will likely reveal more glycans in a glycan fingerprint. MGAT3, mannosylglycoprotein N-acetyl-glucosaminyltransferase 3, specifically recognizes an unmodified GlcNAc residue on the α-3 antenna for substrate recognition [14] and transfers a GlcNAc residue to the β-linked mannose of the trimannosyl core of N-linked oligosaccharides to create a bisecting GlcNAc structure [15], therefore allows us to directly probe some non-galactosylated glycans on IgG.

To perform the experiment, glycans on IgG are first released with PNGase F [16] or Endo S2 [17]. While PNGase F removes glycans entirely from protein backbone, Endo S2 cleaves between the two GlcNAc residues in the chitobiose core of N-glycans and therefore leaves one GlcNAc residue remaining attached to the asparagine residue of the peptide backbone. Our previous study also revealed that Endo S2 prefers short complex N-glycans and has no or little activity on bisecting N-glycans and tri-antennary glycans [18]. For the purpose of glycan identification, the released glycans are then further treated with additional enzymes, such as *C. perfringens* Neuraminidase that specifically removes α-3 and α-6 linked sialic acids [19], B4GalT1 that specifically introduces β-4 linked Gal to non-reducing end GlcNAc residue and forms terminal LN structure [20], α-galactosidase GLA that removes terminal α-linked Gal residues [21], and α-fucosidase FUCA1 that prefers to remove β-6 linked Fuc residues [22]. By comparing glycan fingerprints with and without additional enzymatic treatments, we proved the global existence of core-6 fucosylation and provided evidence of α-Gal epitope containing glycans and paucimannose glycans on NISTmAb and Humira®. The fingerprints also reflect the levels of sialylation and galactosylation of these antibodies.

## Results

### Glycan fingerprinting with Endo S2 for glycan releasing and ST6Gal1 for labeling

In this experiment, glycans of IgG were first released by Endo S2 and then labeled by ST6Gal1 (Fig. 1A) with Cy3-Neu5Ac. Fig. 1B shows the glycan fingerprints of NISTmAb and Humira® in response to Endo S2 inputs. Five bands, designated as band *a*, *b*, *c*, *d*, and *e*, were identified in the fingerprint of NISTmAb (see Table 1 for band designation, mobility, and glycan short names hereafter). Comparing these bands to the Cy3-labeled HA ladder, they migrated to the positions of band 12.5, 10.5, 8.5, 7 and 2.5, respectively. On contrast, three bands (corresponding to band *b*, *c* and *d*) were observed in the fingerprint of Humira®. A fingerprint of thoroughly digested NISTmAb in Fig. 1C further confirmed the bands seen in Fig. 1B. While at the highest input of Endo S2, the heavy chain (HC) of NISTmAb was almost completely deglycosylated, the HC of Humira® was not, suggesting that Humira® may contain some bisecting N-glycans or triantennary glycans that are resistant to Endo S2[18].

**Figure 1.**
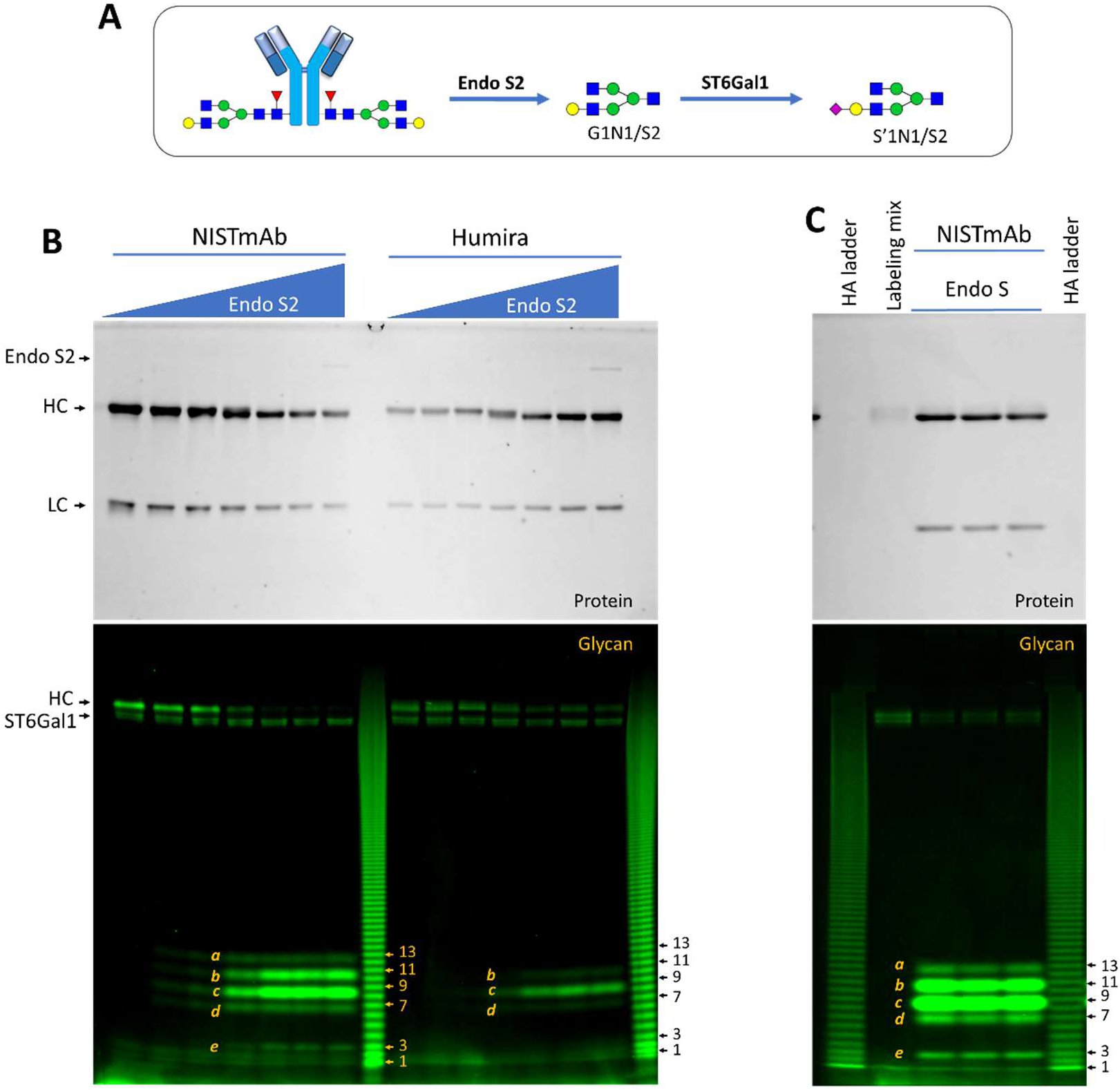
Glycan fingerprinting of NISTmAb and Humira by Endo S2 and ST6Gal1. For rules of short names for glycans, please refer to Note 3 in Table 1. All samples were separated on 17% SDS gel and imaged for both proteins and glycans. **A**) Strategies of N-glycan releasing and labeling. N-glycans are released by Endo S2 under native conditions and labeled by ST6Gal1 with Cy3-Neu5Ac. Short names for the glycans of this study are listed in Table 1. Short names with /S2 [APP]appendix indicate that the glycans were released by Endo S2. **B**) Glycan fingerprints of NISTmAb and Humira in response to Endo S2 inputs. The two antibodies (1 µg each lane) were first treated with variable amounts of Endo S2 (starting from 50 ng with 10-fold serial dilution) at pH 6.5 for 1 hour and then labeled by ST6Gal1 with Cy3-Neu5Ac at 37°C for 2 hours. C) NISTmAb (1 µg) was digested with Endo S2 for overnight and then labeled by ST6Gal1 with Cy3-Neu5Ac for 2 hours at 37°C. An HA ladder was added to monitor the positioning of glycan bands. Each band in the HA ladder is referred as a band number from bottom up (n≥1). Five bands were identifiable in the fingerprint of NISTmAb (designated as *a, b, c, d, e,* at positions close to band 13, 11, 9, 7 and 3, respectively), and four bands corresponding to band *b, c, d, e* were identifiable in the fingerprint of Humira.

When the experiments were repeated with Cy5-Neu5Ac labeling, almost identical results were observed (Supplemental Fig. 1), further validating the method of glycan fingerprinting. Due to that free Cy5 had relatively slow mobility and could interfere with glycan fingerprints, Cy3 was chosen for further study.

### Comparing ST6Gal1 to FUT9 for glycan labeling and PNGase F to Endo S2 for glycan releasing

Due to their unique substrate specificities, only glycans that contain terminal LN are revealed by ST6Gal1 and only glycans without bisecting GlcNAc can be released by Endo S2. To reveal more glycans in a glycan fingerprint, we further explored fingerprinting using FUT9 as a labeling enzyme and PNGase F as a releasing enzyme (Fig. 2A).

**Figure 2.**
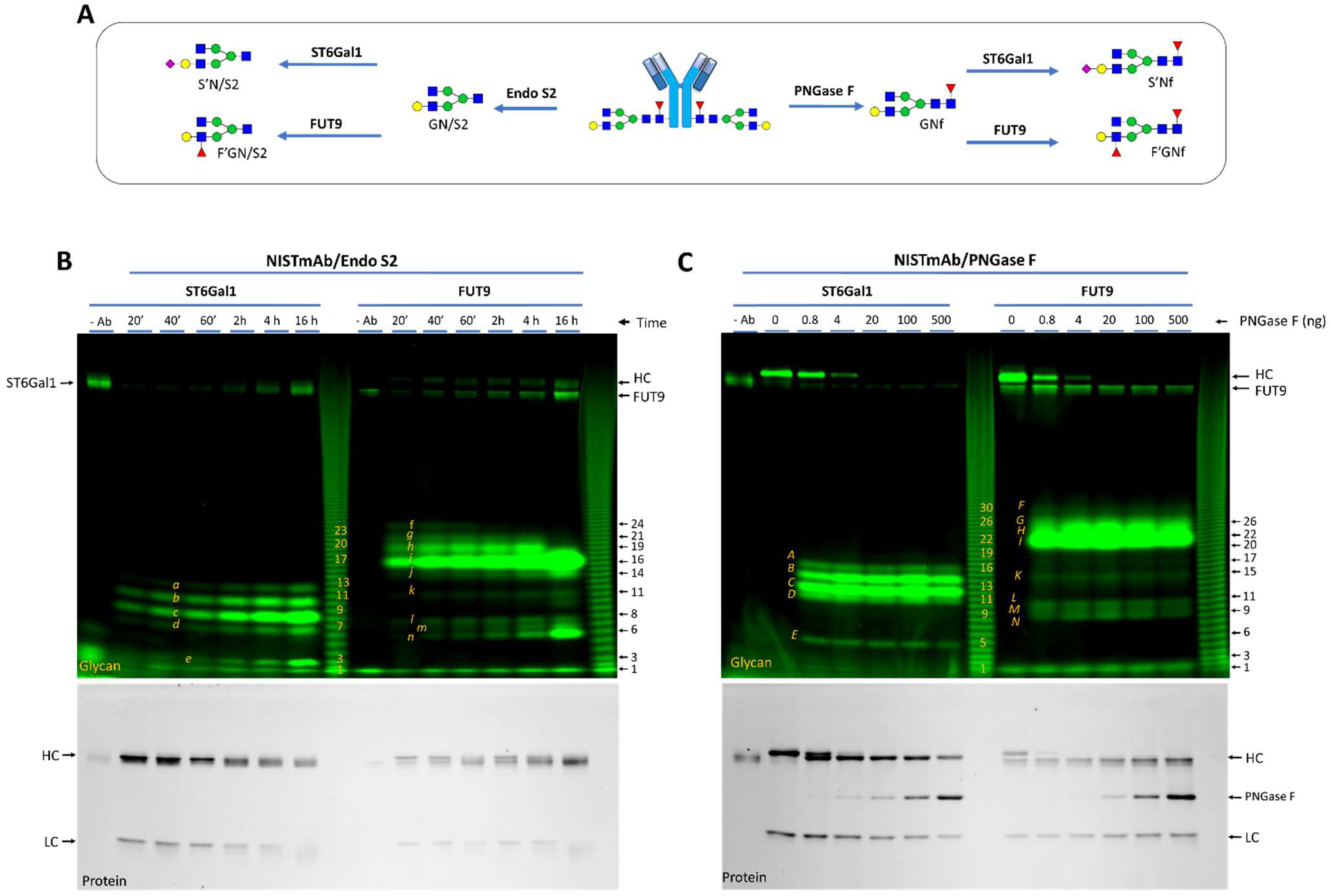
Comparison of Glycan fingerprinting by ST6Gal1 and FUT9. **A**) Strategies for glycan fingerprinting. Glycans are released by Endo S2 or PNGase F and labeled by ST6Gal1 with Cy3-Neu5Ac or by FUT9 with Cy3-Fuc. Note that the glycans released by Endo S2 are short of two sugar residues at their reducing ends. **B**) ST6Gal1 and FUT9 labeling time courses. NISTmAb (1 µg/well) was first treated with Endo S2 (50 ng/well) for one hour then labeled by ST6Gal1 or FUT9 from 20 minutes to 3 hours. **C**) PNGase F dose curve. NISTmAb (1 µg/well) was released by indicated amounts of PNGase F for one hour and then labeled by ST6Gal1 or FUT9 for three hours and one hour, respectively.

A comparison between ST6Gal1 and FUT9 on glycans released from NISTmAb by Endo S2 was performed in Fig. 2B. As expected, five bands were revealed by ST6Gal1, and their intensities increased with time of incubation up to 16 hours. In comparison, nine bands were revealed by FUT9 (band *f* to *n*), with optimal labeling achieved between 40 to 60 minutes of reaction, reflecting the relative fast kinetics of FUT9 [23]. Since glycans recognized by ST6Gal1 should also be labeled by FUT9, the relative intensities of these bands suggest that bands *g*, *h*, *i, j* and *n* correspond to bands *a*, *b*, *c*, *d*, and *e,* respectively, and the extra bands revealed by FUT9 could be due to labeling of glycans with internal LN structures (following experiments further proved this assumption). The intensity of *f* decreased over time and band *g* and *h* greatly reduced at 16 hours of labeling, indicating that these glycans were not stable. For this reason, FUT9 labeling was restricted to 40 to 60 minutes in later experiments. While ST6Gal1 didn’t exhibit obvious labeling, FUT9 showed clear labeling of the HC of Endo S2 treated NISTmAb, indicating that the deglycosylation by Endo S2 was not complete. The remaining glycans on NISTmAb could contain bisecting/tri-antennary GlcNAc residues that were resistant to Endo S2.

To achieve complete deglycosylation, NISTmAb was then treated with PNGase F followed by ST6Gal1 and FUT9 labeling (Fig. 2C). Indeed, PNGase F ≥ 100 ng completely deglycosylated the HC of the antibody, revealed by both ST6Gal1 and FUT9. The glycan fingerprints released by PNGase F had similar band patterning to those released by Endo S2 except that the overall fingerprints were upshifted by two HA band units due to that the PNGase F released glycans contained one additional core GlcNAc residue and possibly one more core Fuc residue than those released by Endo S2 (Fig. 2A). The bands in Fig. 2C were designated as *A*, *B*, *C*, …*L*, *M* and *N,* assuming that there were derived from the same glycans as *a*, *b*, *c*, … *l*, *m*, *n*, respectively.

### Indications of bisecting and tri-antennary glycans on NISTmAb but not Humira®

PNGase F not only releases N-glycans entirely from proteins but also N-glycans that are resistant to Endo S2 such as bisecting N-glycans and highly branched complex glycans. In theory, fingerprints of PNGase F released glycans likely contain more bands than those of Endo S2 released glycans. To this regard, the images of Fig. 2 were re-examined at higher Γ-ratio (a way of changing contrast of an image to purposely increase the exposures of pixels with relatively lower intensity). Indeed, three additional bands were revealed in the fingerprints of PNGase F released glycans (indicated as band α, β, and γ in Supplemental Fig. 2).

A direct comparison between glycan fingerprints of NISTmAb released by PNGase F and Endo S2 further confirmed this finding (Fig. 3). At Γ=2.0, the three additional bands became obvious in the fingerprints of PNGase F released glycans, while no corresponding bands were found in those of Endo S2 released glycans, suggesting that NISTmAb does containing low levels of bisecting and/or tri-antennary glycans. In comparison, these additional bands were not found in the fingerprints of Humira®.

**Figure 3.**
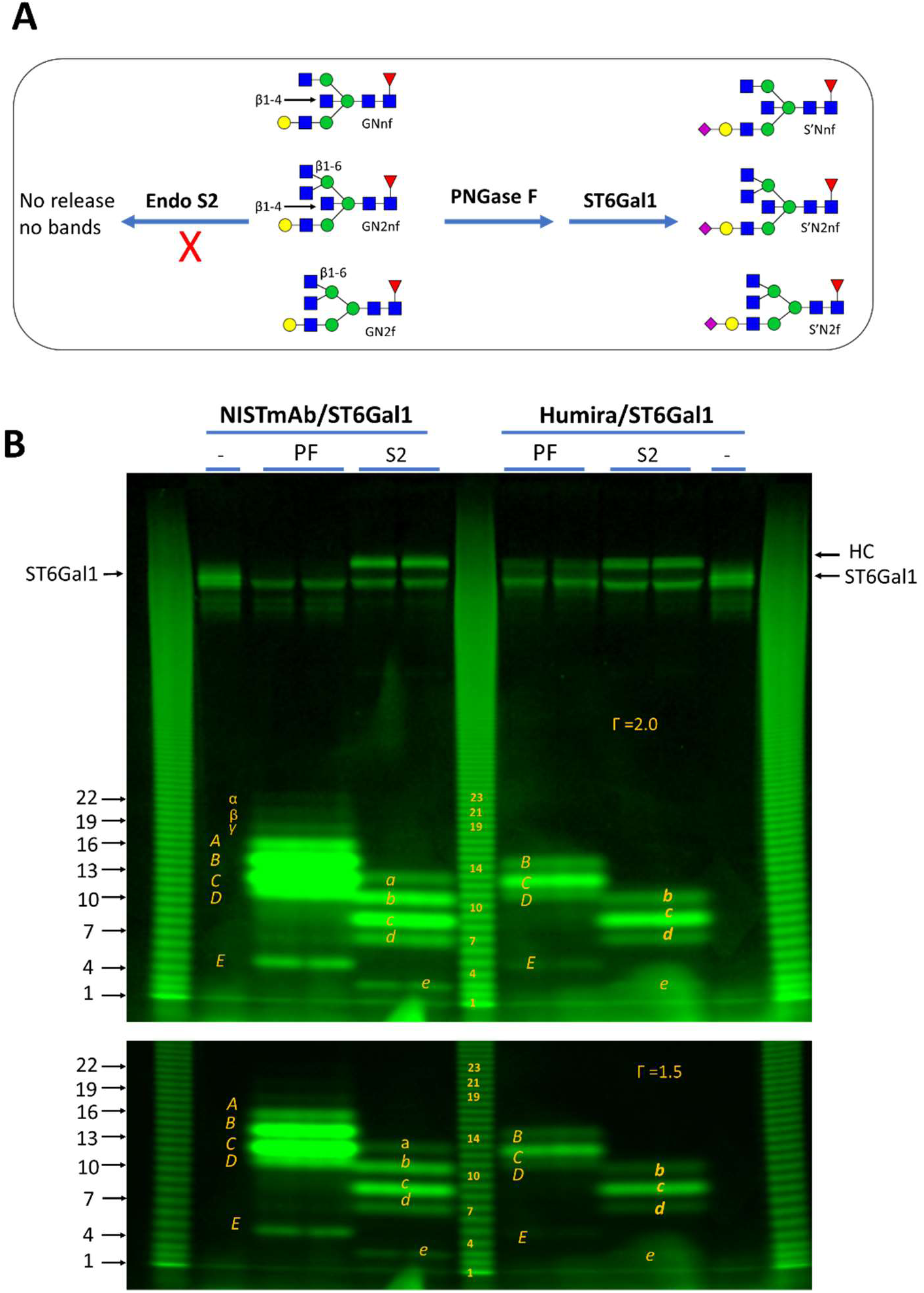
More bands revealed in glycan fingerprints of NISTmAb released by PNGase F than by Endo S2. **A**) Source of glycan variation contributed by PNGase F versus Endo S2. Some variations at the core structures (such as the indicated β1-4 and β1-6 linked GlcNAc residues) render the glycans resistant to Endo S2 digestion and therefore not be revealed in glycan fingerprinting. **B**) Three additional bands (indicated as α, β and γ) from PNGase F (PF) released fingerprints of NISTmAb were observed when Γ was raised to 2.0. Same image with Γ = 1.5 is shown for comparison. No corresponding bands were visible in the fingerprints of Humira. For each reaction, 1 µg of an antibody was digested with PNGase F or Endo S2 (S2) then labeled by ST6Gal1.

### Identify the glycans in the fingerprints of NISTmAb

Considering the limited number of glycan species on NISTmAb, it might be possible to identify those glycans by perturbing the samples with additional enzymatic treatments. To this end, Endo S2 digested NISTmAb was further treated with *C. perfringens* neuraminidase, human α-galactosidase GLA and human B4GalT1 prior to the labeling by ST6Gal1 (Fig. 4A). GLA treatment resulted in the loss of band *a*, suggesting that it contains α-Gal. Since band *a* was labeled by ST6Gal1, it must also have one terminal LN at its non-reducing end. For these considerations, the glycan of band *a* was proposed to be G3. No obvious change was found upon neuraminidase treatment, suggesting that there is no significant level of sialylation. Band *c* was labeled by ST6Gal1 and was completely converted to band *b* by B4GalT1, suggesting that it contains a terminal LN structure and a non-reducing end GlcNAc residue. Considering the limited possible glycans found on NISTmAb, the most likely glycan to satisfy the above requirements for band *c* is GN. Subsequently, band *b* should contain two LN structures and was proposed to be G2. GN and G2 are major glycans on NISTmAb with terminal LN structures [8]. Band *e* responded to B4GalT1 treatment, suggesting the precursor glycan contains a non-reducing end GlcNAc.

**Figure 4.**
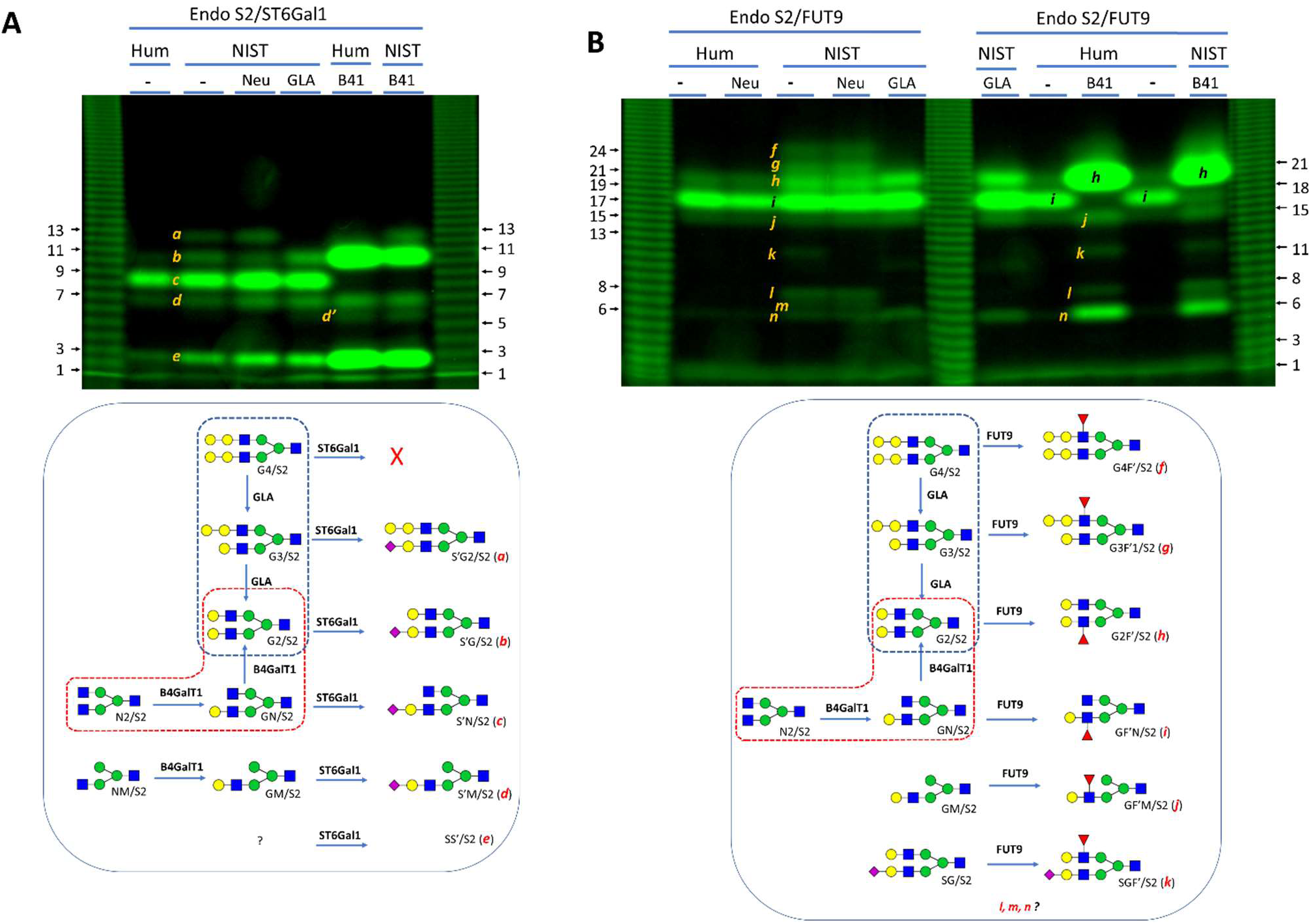
Identifying glycans in the fingerprints of NISTmAb and Humira. For clarity, protein images are omitted and proposed labeling reactions are shown beneath each panel. The glycan conversion routes by α-galactosidase GLA and B4GalT1 are highlighted. **A**) Identify ST6Gal1 labeled glycans. Glycans of NISTmAb and Humira were released by Endo S2 for one hour and then treated with either r*C.p* neuraminidase (Neu) or GLA or B4GalT1 (B41) for 20 minutes prior to labeling by ST6Gal1 for 16 hours. **B**) Identify FUT9 labeled glycans. Glycans of NISTmAb and Humira were released by Endo S2 for one hour and then treated with either Neu or GLA or B41 for 20 minutes prior to labeling by FUT9 for 1 hour.

In Fig. 4B, the same pretreatments were performed but followed by FUT9 labeling. Band *f*, *g, l* and *m* responded to GLA treatment, suggesting that they are α-Gal containing glycans. Considering only one α-Gal containing glycan was identified by ST6Gal1, the additional α-Gal containing glycans identified by FUT9 likely contain internal LN structures that can be recognized only by FUT9. In a separate experiment, it was proved that internal LN structure concealed by α-Gal epitope was only labeled by FUT9 but not ST6Gal1 (Supplemental Fig. 3). It was proposed that band *f* and *g* correspond to G4 and G3 respectively. G4 has two internal LN structures that can only be labeled by FUT9. G3 has both internal and terminal LN structures and can be labeled by both FUT9 and ST6Gal1. The identities of *l* and *m* remained to be studied. Band *k* was the only one responded to neuraminidase treatment and was the weakest band, suggesting that it belongs to a sialylated glycan and that the sialylation level of NISTmAb is very low. Like the conversion of *c* to *b*, *i* was converted to *h* by B4GalT1, suggesting that *i* was derived from GN and *h* was derived from G2. The fact that B4GalT1 pretreatment resulted in far more increase in intensities of *b* and *h* than expected also suggests the presence of N2 that can be converted to G2 via GN by B4GalT1. Like band *e*, band *n* responded to B4GalT1 treatment, but its identity remained to be studied.

### Assessment of core-6 fucosylation on NISTmAb

Core-6 fucosylation is a major factor affecting drug efficacy of therapeutic antibodies [24]. In this experiment, PNGase F treated NISTmAb was further treated with human fucosidase FUCA1 prior to labeling (Fig. 5). FUCA1 treatment resulted in down shifts of all bands in PNGase F released fingerprints regardless they were labeled by ST6Gal1 or FUT9, suggesting that all labeled glycans contain core-6 fucose. The possibility of the fucose at other location was unlikely as fucosylation at LN structures would prevent FUT9 and ST6Gal1 labeling.

**Figure 5.**
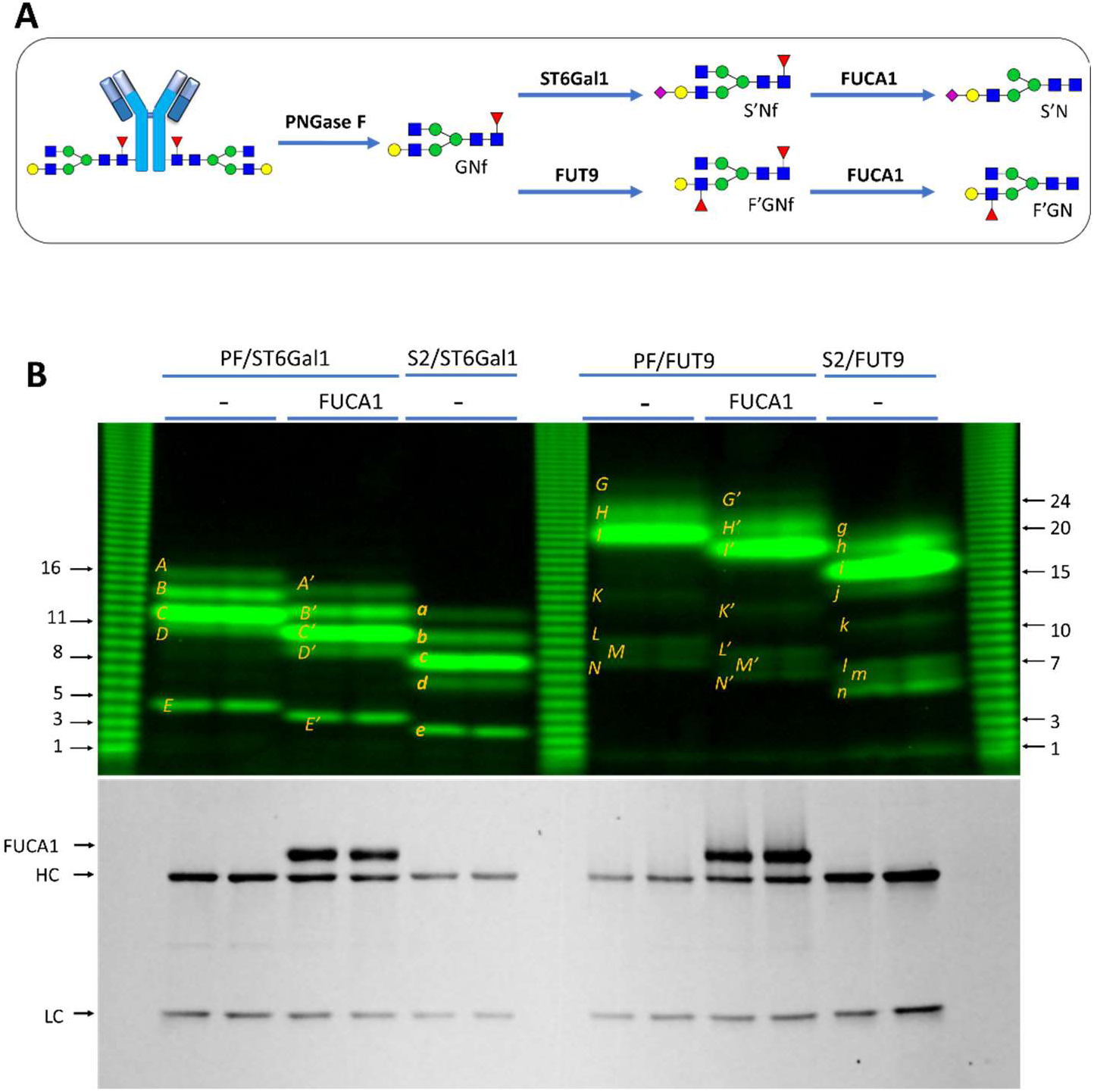
Probing fucosylation on NISTmAb. **A**) Strategy. **B**) Fingerprints along with protein image. NISTmAb (1 ug/well) was first digested by PNGase F (PF) or Endo S2 (S2) alone. The digestions were then labeled by ST6Gal1 for 3 hours or FUT9 for one hour. Portion of the PF-released ST6Gal-labeled glycans were further treated with FUCA1. All samples were separated on 17% gel. Protein image is also shown.

### Detecting non-galactosylated glycans by MGAT3

B4GalT1 treatment can convert some non-galactosylated glycans (such as N2) to galactosylated glycans for ST6Gal1 and FUT9 detection (Fig. 4). However, it is more desirable if these glycans can be directly detected. To this end, Endo S2 released glycans from NISTmAb and Humira® were directly labeled by MGAT3 with UDP-AF488-GlcNAc (Fig. 6). The MGAT3 labeled fingerprint of NISTmAb contained two relatively weak bands (*o* and *p*) and that of Humira® contained one strong band (*p*). Based on the substrate specificity of MGAT3, the limited number of known glycans present on NISTmAb and Humira® and their difference on mobility, it is postulated that *p* was derived from N2 and *o* was derived from M2N. Fig. 6 also suggests that N2 is far more abundant in Humira® than NISTmAb.

**Fig. 6.**
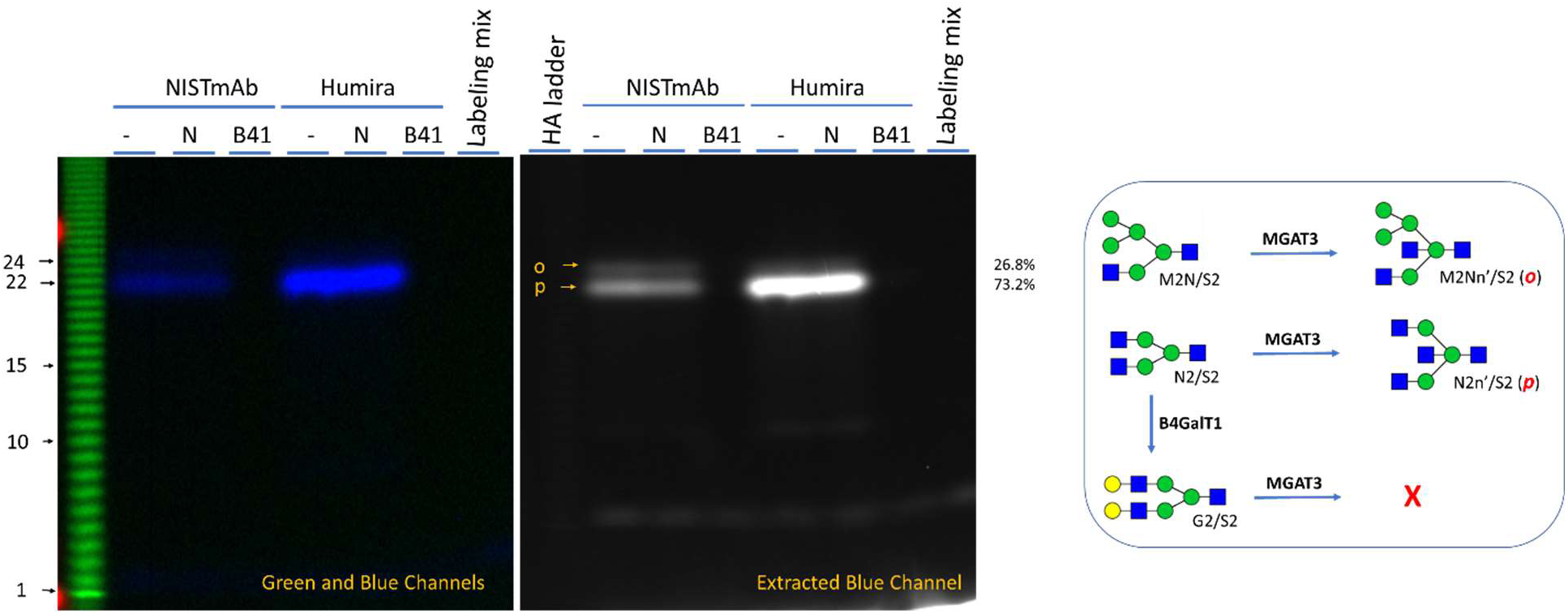
Glycan fingerprinting of NISTmAb and Humira by MGAT3 with AF488-GlcNAc Glycans of all samples were released with Endo S2, labeled by MGAT3 with AF488-GlcNAc and separated on 17% gel. Cy3-labeled HA ladder was run along the samples. Proposed glycans and labeling mechanisms are shown beneath the image. The left side is the composite image of both green and blue channels. For better viewing the signals, the right side shows the extracted image from the blue channel only. Glycans of NISTmAb and Humira were released by Endo S2 for one hour and then treated with either r*C.p* neuraminidase (N) or B4GalT1 (B41) for 20 minutes prior to labeling by MGAT3 for 16 hours. B41 treatment abolished the labeling. While both band *o* and *p* are barely visible from NISTmAb, a much stronger band *p* is visible from Humira, suggesting that Humira contains primarily N2 glycan.

### Detecting paucimannose glycans by FUT8

Band *o* in Fig. 6 suggests the presence of paucimannose glycans in NISTmAb. Paucimannose glycans are important indicators for glycan maturation and have been reported on NISTmAb [8]. To detect these glycans, samples were either directly labeled with Cy5-Fuc by FUT8 or after pretreatment with MGAT1 and/or FUCA1 (Fig. 7). MGAT1 [25] adds a β1-2 linked GlcNAc residue to the α1-3 linked Man residue of the paucimannose glycans so that these glycans become the substates for FUT8 [26]. SF21 insect cell expressed SARS2 RBD protein that contained both M2 and M3 was used as a control [9]. Three major bands (*R*, S and *T*) were observed. *R* and *T* had the same mobility as the bands observed in the control, suggesting that that they are M2Nf’ and MNf’, respectively. Band *T* was observed on both NISTmAb and Humira ® without prior pretreatment and its levels were increased by FUCA1 treatment, suggesting the presence of both MN and MNf in both antibodies. Band *R* was observed on both antibodies only after MGAT1 treatment and its levels were not increased significantly by FUCA1 treatment, suggesting the existence of relatively high levels of M3 but not M2Nf.

**Fig. 7.**
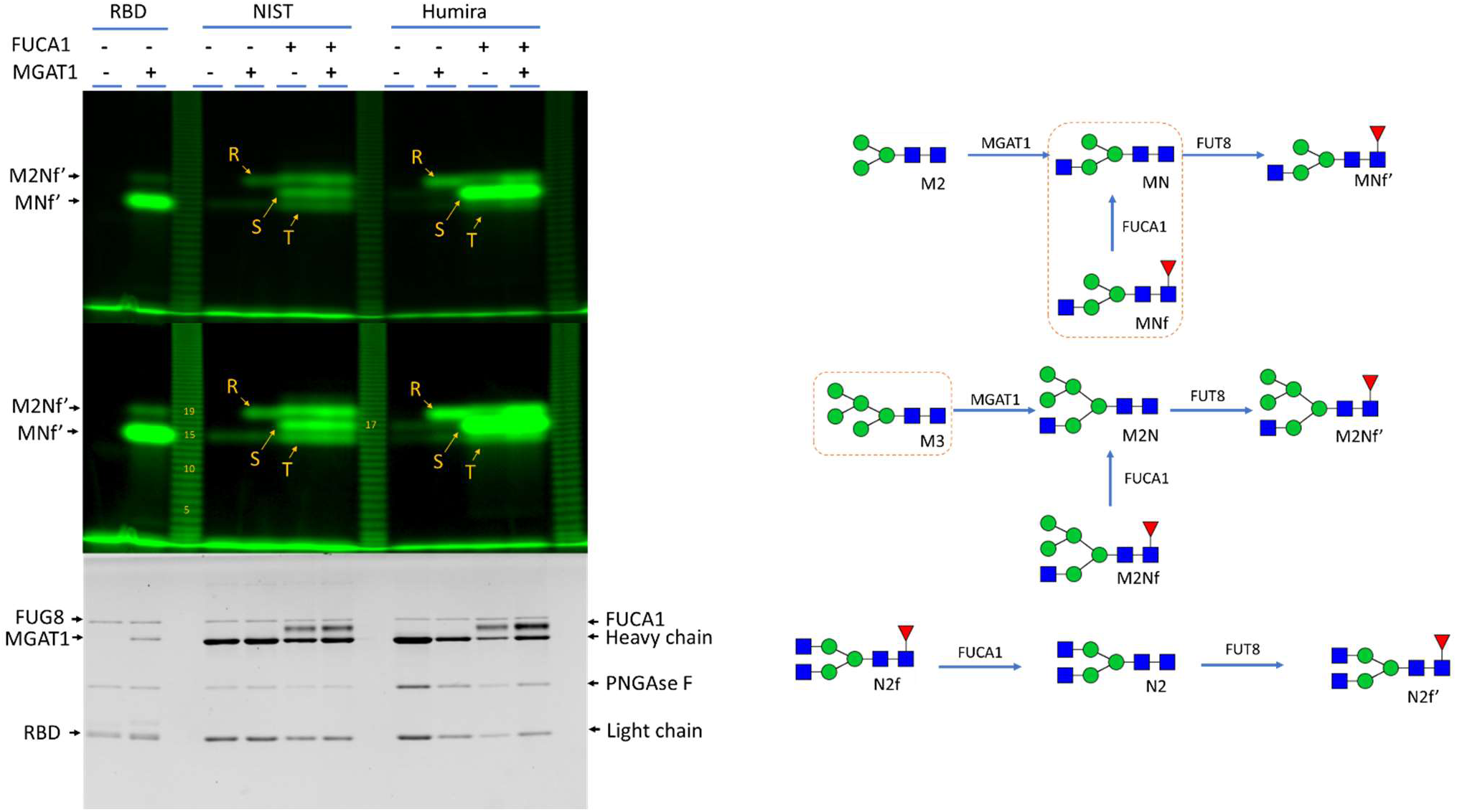
Detecting paucimannose glycans of NISTmAb and Humira. Glycans were released with PNGase F, briefly treated with B4GalT1, heat inactivated, then treated with MGAT1 and/or FUCA1, heat inactivated again and finally labeled with Cy3-Fuc by FUT8. SARS2 RBD protein (expressed in insect cells) that contains M2 and M3 glycans served as control. All samples were separated on 17% gel. Cy3-labeled HA ladder was run along the samples. The gel was imaged for glycans (top two panels with different contrasts) and proteins (lower panel). Proposed glycan labeling mechanisms are shown beneath the images.

Band *S* moved between *R* and *T* and was observed on both antibodies but not on control and its intensities were dramatically increased by FUCA1 treatment, suggesting that it originated from N2f. The sensitivity of band *S* to FUCA1 treatment is consistent to Fig.5. Besides, the intensity difference of band *S* on Humira ® and NISTmAb was consistent to that of band *p* in Fig. 6, further confirming that N2f is much more abundant on Humira®.

## Discussion

In this report, using glycan fingerprinting we studied glycosylation of NISTmAb and Humira® with different combinations of glycan releasing, modifying and labelling enzymes. Overall, 15 unique glycan bands in six groups were identified in the fingerprints of NISTmAb (Table 1). Group I was resistant to Endo S2 digestion, had slowest mobility among all ST6Gal1 labeled bands, and were only revealed at extremely high gamma rating (2.0), suggesting that they are rare structures with bisecting and/or β1,6-linked GlcNAc residues. Group II was sensitive to α-Galactosidase, therefore must contain α-Gal epitopes. Group III was released by both PNGase F and Endo S2 and revealed by both ST6Gal1 and FUT9. Group IV was revealed by MGAT3, suggesting that they contain unmodified α3 arm GlcNAc residue. Group V were only revealed by FUT9 and sensitive to fucosidase, suggesting that all three bands contain internal LN structures and core fucose. Group VI had fasted mobility, was released by both PNGase F and Endo S2, labeled by both ST6Gal1 and FUT9, and was sensitive to fucosidase, suggesting that the glycan is likely negatively charged [9], contains terminal LN and core fucose. Group II, III and IV are relatively abundant and their structures were proposed based on the information available. Group V and VI were relatively rare and had much faster mobility than other groups, and their structures are beyond speculation.

The overall glycosylation on Humira® is simpler, with only group III, IV and VI glycans detected. The most noticeable thing of Humira® is that no α-Gal was detected. Although no group I glycans were directly detected from Humira® (Fig. 3), the antibody did contain Endo S2 resistant glycans (Fig. 1 and Fig.3), indicating that it contains some other type of complex glycans.

From a technical perspective, our methods are characterized by their convenience, sensitivity, quantitative accuracy, and consistency. The assay procedures consist of glycan releasing, modification, and labeling, all under physiological conditions, and can be finished within 2 days with hands on time less than 2 hours. The labeled samples are separated with electrophoresis and visualized with a fluorescent imager. When the samples are run together with a HA glycan ladder, all resolved bands can be assigned with unique HA ladder numbers, therefore allowing data obtained from different experiments comparable. A glycan fingerprint can be generated with as little as sub microgram of an antibody and the band intensities are directly related to the abundance of glycans. Consistency is evidenced on the band patterning revealed in different experiments. Additionally, the data generated are visual and relatively straightforward to interpret.

Another advantage of the assay is that it allows monitoring released glycans as well as glycans that remain attached to the antibodies simultaneously, whereas most current methods study released glycans without knowing if all glycans have been released from antibodies.

The limitation of the assay is due to the restricted substrate specificity of labeling enzymes. Each enzyme has its unique substrate preference. Glycans that are not substrate to a labeling enzyme will not be revealed in a glycan fingerprint. However, this issue can be mitigated by using combinations of different enzymes. In this report, glycans with terminal LN were labeled by ST6Gal1, both terminal and internal LN structures were labeled by FUT9, glycans without LN structure maybe labeled by MGAT3, and glycans without core –fucose could be labeled by FUT8. In a sense, these enzymes are complementary to each other. In addition, glycans of one type may be converted to another type for detection. For examples, glycans without LN may be detected by pre-installment of the structure with B4GalT1 (Fig. 4), and oligomannose can be converted to the substrate of FUT8 for detection [9]. With the wide availability of glycosyltansferases and glycosidases, more and more glycans can be labeled and detected using our methods.

## Material and methods

NISTmAb was directly obtained from the National Institute of Standards and Technology. Humira® was directly from Abbott Laboratories. Recombinant human FUCA1, GLA, ST6Gal1, FUT9, B4GalT1, MGAT3, *S. pyogenes* Endo S2, *C. perfringens* neuraminidase, *F. meningosepticum* PNGase (PNGase F), CMP-Cy5-Neu5Ac, CMP-Cy3-Neu5Ac, GDP-Cy3-Fuc, UDP-AlexaFluor488-GlcNAc, Cy3-lableled HA ladder were from Bio-Techne. All other chemicals from Sigma Aldrich.

### Releasing N-Glycans from the NISTmAb and Humira® and additional enzymatic treatment

The following are general protocols. Individual experiments vary depending on the purposes. To release N-glycans, 10 μg of an antibody was mixed with variable amounts of PNGase F or Endo S2 and supplemented with 25 mM MES pH 6.5, 150 mM NaCl to 20 μL and then incubated at 37° C for at least 30 minutes. For further desialylation treatment, an additional 0.2 μg *C. perfringens* neuraminidase was added into the reaction mixture and incubate for additional 30 minutes. For GLA and FUCA1 treatment, the buffer pH was adjusted to 4.5 with citric buffer and 2 µg of GLA or FUCA1 was added to the reaction and the reaction was incubated for another 30 minutes. For B4GalT1 treatment, 0.2 µl of the enzyme, 1 mM of UDP-Gal and 10 mM of MnCl_2_ was added to the reaction and the reaction was further incubated for another 30 minutes. The above reactions were then heated at 95° C for two minutes to inactivate the enzymes.

### Labeling released glycans from NISTmAb and Humira®

The following are general protocols. Individual experiments vary depending on the purposes. For labeling with ST6Gal1, 4 µg of treated antibodies was mix with 0.2 μg of ST6Gal1, 0.1 nmol of CMP-Cy3-Neu5Ac (or CMP-Cy5-Neu5Ac) and supplemented with labeling buffer (25 mM Tris pH 7.5, 10 mM MnCl) to 20 μL. For labeling with FUT9, 4 µg of treated antibodies was mixed with 1 μg of FUT9, 0.1 nmol of GDP-Cy3-Fuc and supplemented with labeling buffer (25 mM Tris pH 7.5, 10 mM MnCl) to 20 μL. For labeling with MGAT3, 4 µg of treated antibodies was mixed with 1 μg of MGAT3, 0.1 nmol of UDP-AlexaFluor488-GlcNAc and supplemented with labeling buffer (25 mM Tris pH 7.5, 10 mM MnCl) to 20 μL. The labeling reactions were then incubated at 37° C for 60 minutes or overnight at room temperature.

### Glycan Electrophoresis and Imaging

One μg of labeled antibody was loaded to each well of a 17% sodium dodecyl sulfate– polyacrylamide gel and the gel was run at 20 volts/cm till dye front reached the end of the gel. After separation, the gel was first imaged for glycans using a FluorChem M imager (ProteinSimple, Bio-techne) under red or green or blue channel and then imaged with traditional methods such as silver staining or trichloroethanol (TCE) staining for protein contents.

## Author contributions

ZW performed the experiments and wrote the main manuscript and prepared the figures.

## Acknowledgements

The author would like to thank his former colleagues at Bio-techne who provided the reagents for this study.

## Data availability statement

All data generated or analyzed during this study are included in this published article (and its Supplementary Information files).

---

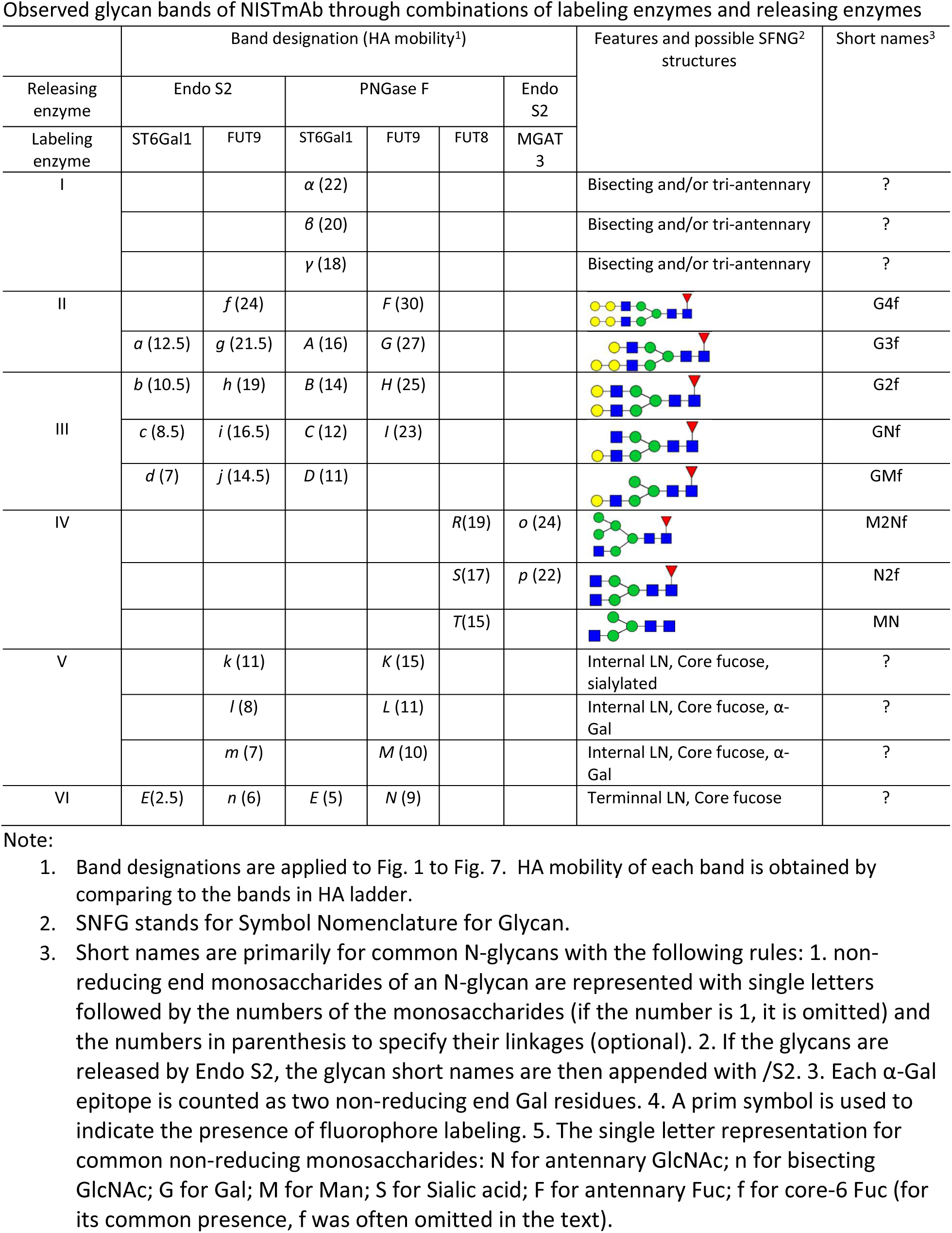
Table.

| Releasing enzyme | Band designation (HA mobility<sup>1</sup>) |  |  |  |  |  | Features and possible SFNG<sup>2</sup> structures | Short names<sup>3</sup> |
| --- | --- | --- | --- | --- | --- | --- | --- | --- |
|  | Endo S2 |  | PNGase F |  |  | Endo S2 |  |  |
| Labeling enzyme | ST6Gal1 | FUT9 | ST6Gal1 | FUT9 | FUT8 | MGAT 3 |  |  |
| I | | | $\alpha$ (22) | | | | Bisecting and/or tri-antennary | ? |
| | | | $\beta$ (20) | | | | Bisecting and/or tri-antennary | ? |
| | | | $\gamma$ (18) | | | | Bisecting and/or tri-antennary | ? |
| II |  | *f* (24) |  | *F* (30) |  |  |  | G4f |
|  | *a* (12.5) | *g* (21.5) | *A* (16) | *G* (27) |  |  |  | G3f |
| III | *b* (10.5) | *h* (19) | *B* (14) | *H* (25) |  |  |  | G2f |
|  | *c* (8.5) | *i* (16.5) | *C* (12) | *I* (23) |  |  |  | GNf |
|  | *d* (7) | *j* (14.5) | *D* (11) |  |  |  |  | GMf |
| IV |  |  |  |  | *R*(19) | *o* (24) |  | M2Nf |
|  |  |  |  |  | *S*(17) | *p* (22) |  | N2f |
|  |  |  |  |  | *T*(15) |  |  | MN |
| V |  | *k* (11) |  | *K* (15) |  |  | Internal LN, Core fucose, sialylated | ? |
| | | *l* (8) | | *L* (11) | | | Internal LN, Core fucose, α-Gal | ? |
| | | *m* (7) | | *M* (10) | | | Internal LN, Core fucose, α-Gal | ? |
| VI | *E*(2.5) | *n* (6) | *E* (5) | *N* (9) |  |  | Terminal LN, Core fucose | ? |

Note:

**Supplemental Figure 1.**
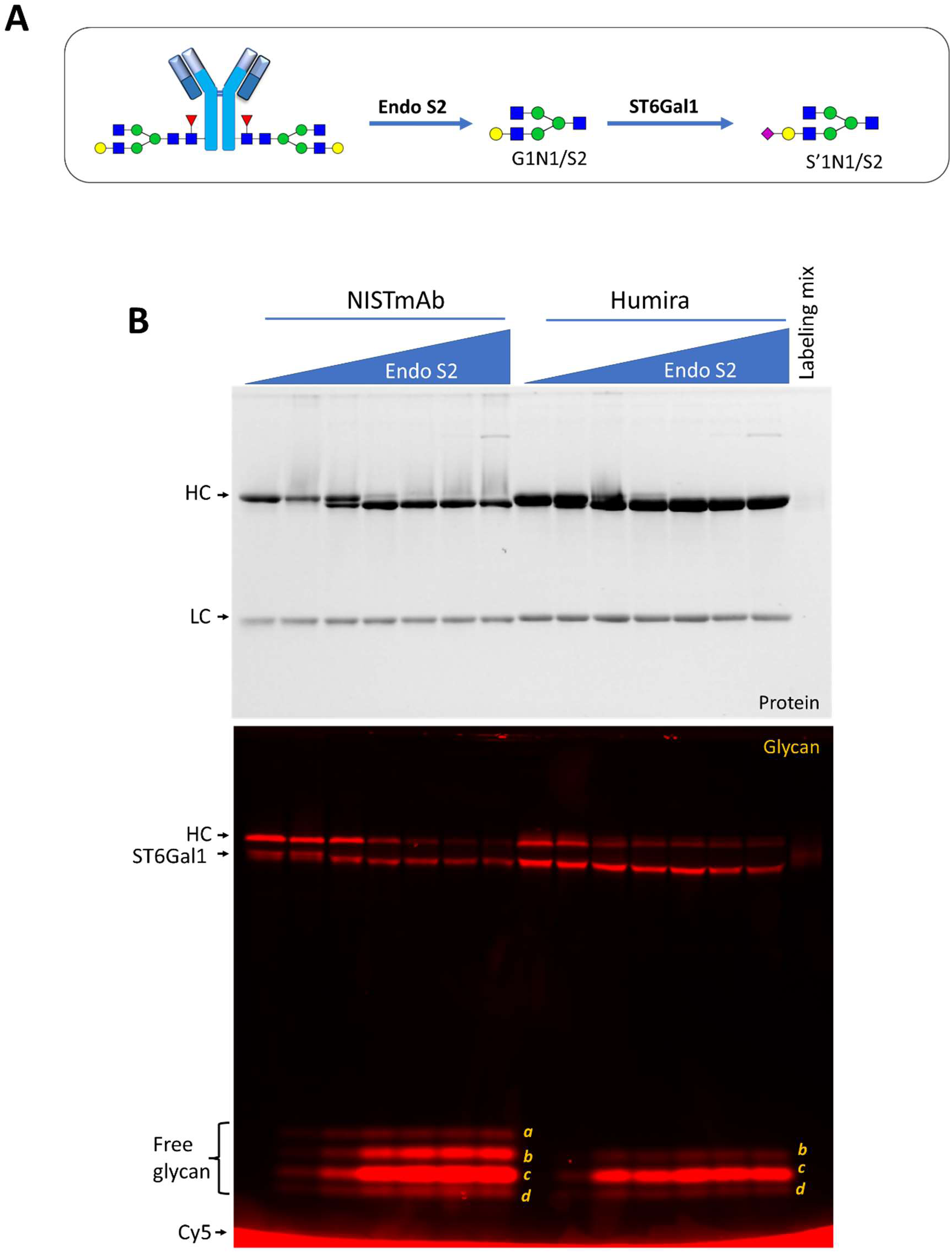
Glycan fingerprinting of NISTmAb and Humira by Endo S2 and ST6Gal1 with Cy5-Neu5Ac. All samples were labeled by ST6Gal1 and separated on 17% SDS gel and imaged for both proteins (upper panel) and labeled glycans (lower panel). To reveal the weak bands, the γ rating was increased to 1.5. A) Strategies of N-glycan releasing by Endo S2 and labeling by ST6Gal1. B) NISTmAb (4 µg) and Humira (8 µg) were first treated with Endo S2 (starting from 200 ng with 5-fold serial dilution) at pH 6.5 for 1 hour and then labeled by ST6Gal1 with Cy5-Neu5Ac at 37°C for 2 hours. Four bands likely corresponding to band *a, b, c*, and *d* and three bands likely corresponding to band *b, c* and *d* of Fig. 1B can be identified in the glycan fingerprints of NISTmAb and Humira, respectively.

**Supplemental Figure 2.**
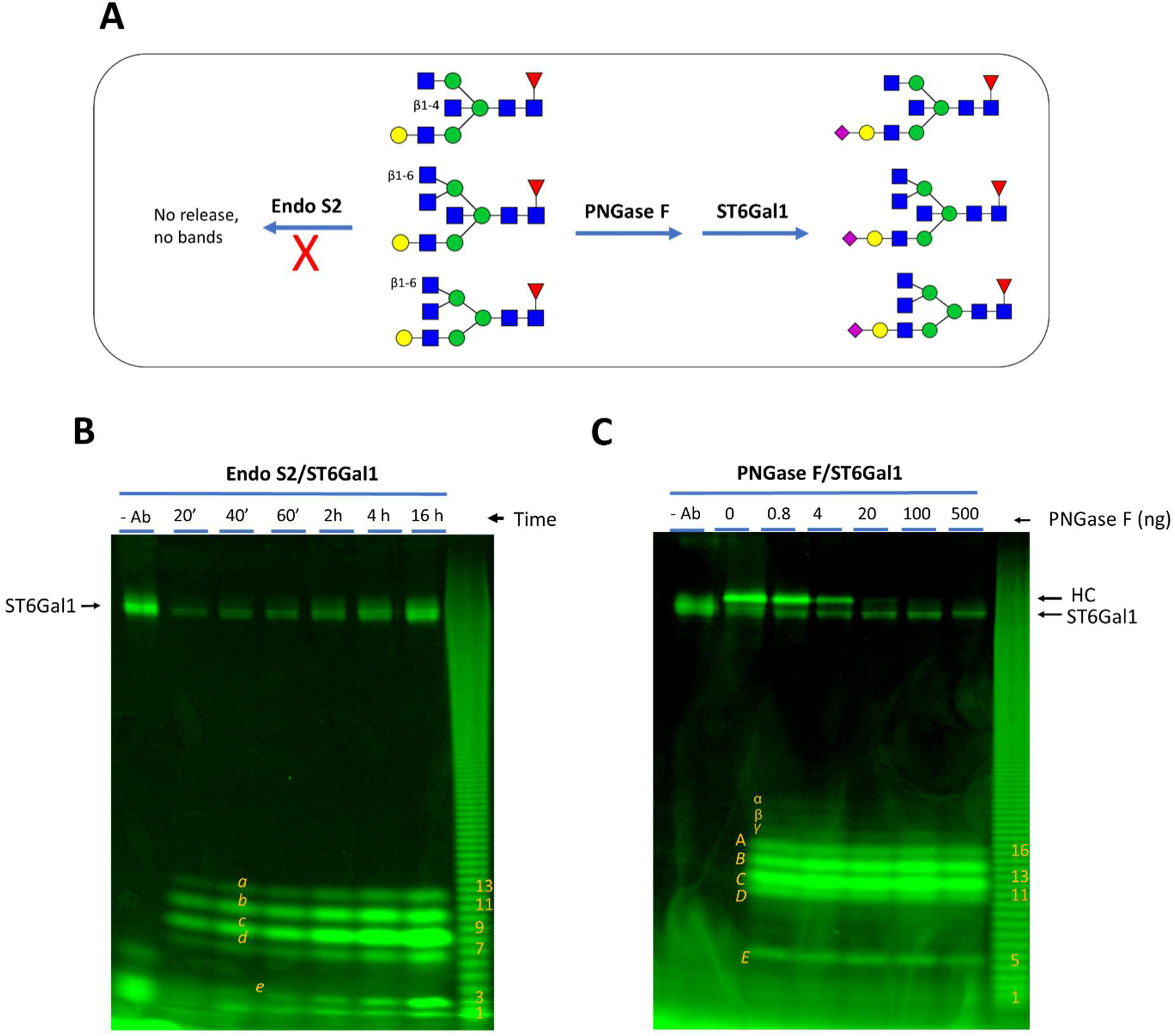
Additional bands were revealed by raising the gamma rating to 2.0 for the images in Figure 2. **A**) Source of glycan variation contributed by PNGase F versus Endo S2. Some variations at the core structures (such as the indicated β1-4 and β1-6 linked GlcNAc residues) render the glycans resistant to Endo S2 digestion and therefore not be revealed in glycan fingerprinting. **B**) The image of Endo S2 released glycans labeled by ST6Gal1. **C**) The image of PNGase F released glycans labeled by ST6Gal1. Three additional bands (indicated as α, β and γ) in B are visible. These bands are believed to be caused by variations of the core structures such as the introduction of the β1-6 linked GlcNAc by MGAT5, or the introduction of the β1-4 linked GlcNAc by MGAT3.

**Supplemental Fig. 3.**
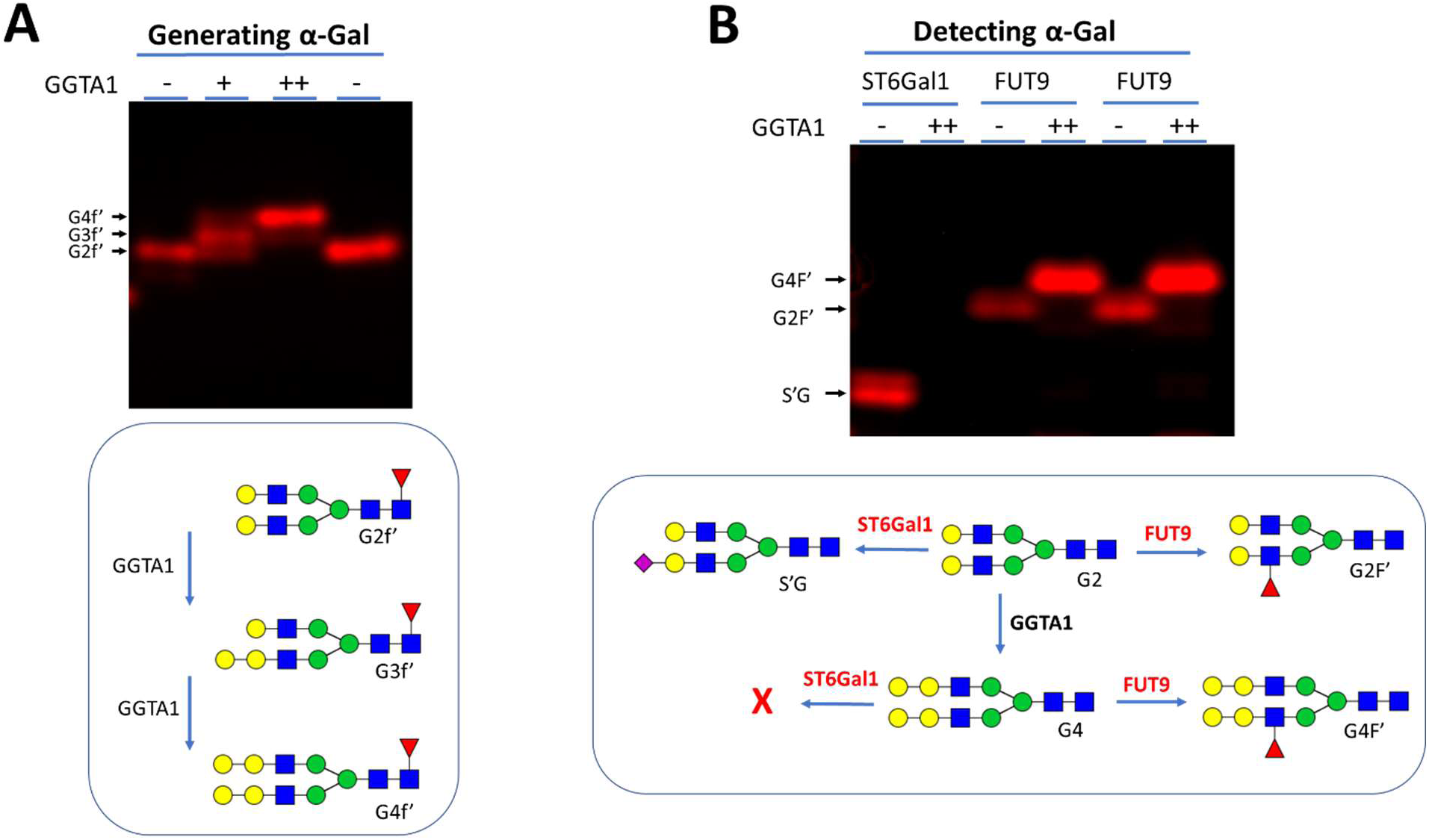
Detecting α-Gal epitope by ST6Gal1 and rhFUT9. Proposed molecular mechanisms are shown below each image. **A**) Generating α-Gal on labeled G2f’. α-Gal was generated on G2f’ by modifying with recombinant mouse GGTA1. Gel shift caused by the treatment suggests the formation of either one (G3f’) or two α-Gal epitopes (G4f’). **B**) In a parallel experiment, G2 was modified GGTA1 exhaustively then labeled by ST6Gal1 and FUT9 with Cy5-Neu5Ac and Cy5-Fuc, respectively. GGTA1 modification abolished ST6Gal1 labeling, suggesting that ST6Gal1 do not recognize α-Gal capped lactosamine structure. On contrary, GGTA1 modified glycans were strongly labeled by FUT9, suggesting that FUT9 recognize α-Gal capped lactosamine structures.

